# The circadian dynamics of small nucleolar RNA

**DOI:** 10.1101/102533

**Authors:** Stuart Aitken, Colin A. Semple

## Abstract

The circadian regulation of gene expression allows plants and animals to anticipate predictable environmental changes. While the influence of the circadian clock has recently been shown to extend to ribosome biogenesis, the dynamics and regulation of the many small nucleolar RNA that are required in pre-ribosomal RNA folding and modification are unknown. Using a novel computational method, we show that 18S and 28S pre-rRNA are subject to circadian regulation in a nuclear RNA sequencing time course. A population of snoRNA with circadian expression is identified that is functionally associated with rRNA modification. More generally, we find the abundance of snoRNA known to modify 18S and 28S to be inversely correlated with the abundance of their target. Cyclic patterns in the expression of a number of snoRNA indicate a coordination with rRNA maturation, potentially through an upregulation in their biogenesis, or their release from mature rRNA at the end of the previous cycle of rRNA maturation, in antiphase with the diurnal peak in pre-rRNA. Few cyclic snoRNA have cyclic host genes, indicating the action of regulatory mechanisms in addition to transcriptional activation of the host gene. For highly-expressed independently transcribed snoRNA, we find a characteristic RNA polymerase II and H3K4me3 signature that correlates with mean snoRNA expression over the day.

## 1 Introduction

Circadian rhythms in animal physiology and metabolism anticipate predictable diurnal variations in the environment [1, 2]. In mammals, the master clock is located in the hypothalamic suprachiasmatic nucleus (SCN) region of the brain. Cells and peripheral organs have autonomous oscillators coordinated with the SCN through hormonal signals [3]. The molecular basis of these cycles is well understood: The Clock:Bmal1 heterodimer activates the transcription of Per and Cry genes, these proteins then repress Clock and Bmal1 transcription [2, 3]. Despite Clock:Bmal1 binding its targets in a narrow time window, 6 hrs after dawn in mouse, the targeted genes peak in expression at varying times [4]. Recent genome-wide sequencing studies of nascent and mature mRNA have shown that rhythmic premRNA transcription is not necessarily followed by rhythmic mRNA levels, and that rhythms in mRNA expression are observed in genes lacking rhythmic transcription [5]. In addition, oscillations in protein levels and phosphorylation states give further evidence for circadian regulation operating at all levels from transcription to translation, splicing and the maintenance of transcript stability [6, 7].

Ribosome biogenesis has also been shown to be influenced by the circadian clock through the transcription of translation initiation factors, ribosomal proteins and ribosomal RNAs [8]. For example, in mouse, Rps18 and Rpl30 mRNA expression in the polysomal RNA fraction peaks at 14-22 hrs after dawn [8]. The 45S pre-rRNA peaks around the middle of the day and is synchronised with ribosomal proteins potentially through UBF1 binding [8]. However, little is known about the dynamics of the complex process of rRNA biogenesis across the circadian cycle, or the regulation of the many small nucleolar RNAs (snoRNAs) that support the folding and modification of rRNA precursors.

Small nucleolar RNA are short noncoding RNAs with a conserved role in ribosome biogenesis. SnoRNAs are found in both eukaryotes and archaea indicating an ancient origin [9]. In mammals, many of the currently characterised snoRNA are located in the introns of protein-coding genes, from which they are processed after splicing and debranching of the intron lariat [10]. SnoRNA are also found embedded within annotated noncoding host genes (named processed transcripts in current genome bio-type annotations), lincRNAs, and in nongenic regions. Two classes of snoRNA have been defined, box C/D and box H/ACA, guiding the methylation of rRNA and its pseudouridylation respectively. Box C/D snoRNAs have an additional role in the cleavage and folding of pre-rRNA. However, numerous orphan snoRNAs outside of these two classes are also known to exist in mammalian genomes, and a variety of novel functional roles for snoRNAs have also emerged. Suggested non canonical functions of snoRNA include the cross-modification of other snoRNA, binding to other ncRNAs (e.g. 7SK), the editing and splicing of mRNA, association with accessible chromatin and as precursors for miRNA [10]. A potential role for snoRNA in circadian metabolism in mouse and human has also been suggested [11], and snoRNA host genes in Drosophila have been shown to oscillate [12] but, to date, the extent of circadian dynamics across the diversity of mammalian snoRNA transcripts is unknown.

Here we take advantage of unique nascent (poly A-) and mature (poly A+) RNA sequencing data sets to explore the expression dynamics of ribosome biogenesis in mouse liver. We show that nascent RNA-seq data is a rich resource that reveals both snoRNA and pre-rRNA dynamics, and using a new approach to detect periodic expression we reveal novel subpopulations of circadian snoRNA and a distinct subpopulation with with time-varying expression greatly in excess of their host genes. Additional data on chromatin state gives further novel insights into snoRNA biogenesis. Overall these data suggest that snoRNAs regulated with circadian periodicity are tightly integrated with ribosome biogenesis in mammalian cells.

## 2 Results

We quantify the remarkable variations in snoRNA, host mRNA and rRNA abundance, and explore their interrelationships, in next generation sequencing data generated across the circadian cycle in mouse liver [5, 13]. Cyclical variations in microRNA expression across the circadian cycle have been noted [14, 15], as have such variations in lincRNA [5], but to date such changes in snoRNA and rRNA have not been revealed. We adopt the current mouse assembly (Ensembl GRCm38) and annotation (84) for coding and noncoding genes. Approximately 1500 snoRNA genes are included, many from RFAM computational predictions (which have had a controversial status [16]). Thus we explore the current catalog of snoRNA gene expression in a diverse range of sequencing data from mouse.

### 2.1 SnoRNA are a major constituent of nascent sequencing data

Nascent sequencing captures nuclear RNA prior to the formation of the 3’ end [5]: The protocol depleted polyadenylated RNA but rRNA was not removed. Thus we found the nascent sequencing data contained reads mapping to most RNA species, including those that are not polyadenylated in their mature form such as snoRNA and rRNA. Two biological samples were obtained at six time points from Zeitgeber Time 0 (ZT0, dawn) to ZT20 (20 hours after dawn) from 12 different mice (see [5] for details).

The abundance of RNA transcripts was quantified in TPM using the Kallisto pseudo-alignment technique [17]. This requires the set of transcripts of interest to be compiled (Ensembl GRCm38) to which we added the 5.8S, 18S and 28S pre-rRNA sequences (snOPY database [18]). The large proportions of snoRNA and rRNA species, and their variation over the day in nascent seq data were unexpected but readily apparent (Fig 1a). It was evident that mRNA constituted between 16% and 27% of the RNA in nascent sequencing across the day, with rRNA, snoRNA and snRNA all accounting for at least 15% of sequenced RNA. In contrast, mRNA constituted over 94% of RNA abundance in conventional RNA sequencing data (Fig 1b).

### 2.2 SnoRNA hosted by protein-coding genes and nongenic snoRNA are extensively expressed in mouse liver

There is a considerable discrepancy between the number of snoRNA genes curated in the literature and the number annotated as snoRNA in Ensembl, based on a computational prediction protocol. The number of snoRNA genes in mammals has been estimated as 216 (*H. sapiens*) [19], which is only a fraction of the 1484 snoRNA annotated in the mouse assembly GRCm38 (mm10). To address whether these genes are expressed in mouse liver, for each of the 12 sequencing datasets, we selected a lower threshold of TPM expression as the first quartile (0.7-1.6 TPM) and considered all genes with expression above this threshold in any dataset to be expressed. As an additional test, we required at least a single uniquely-mapping read per transcript to call a snoRNA identifiable.

**Figure 1:**
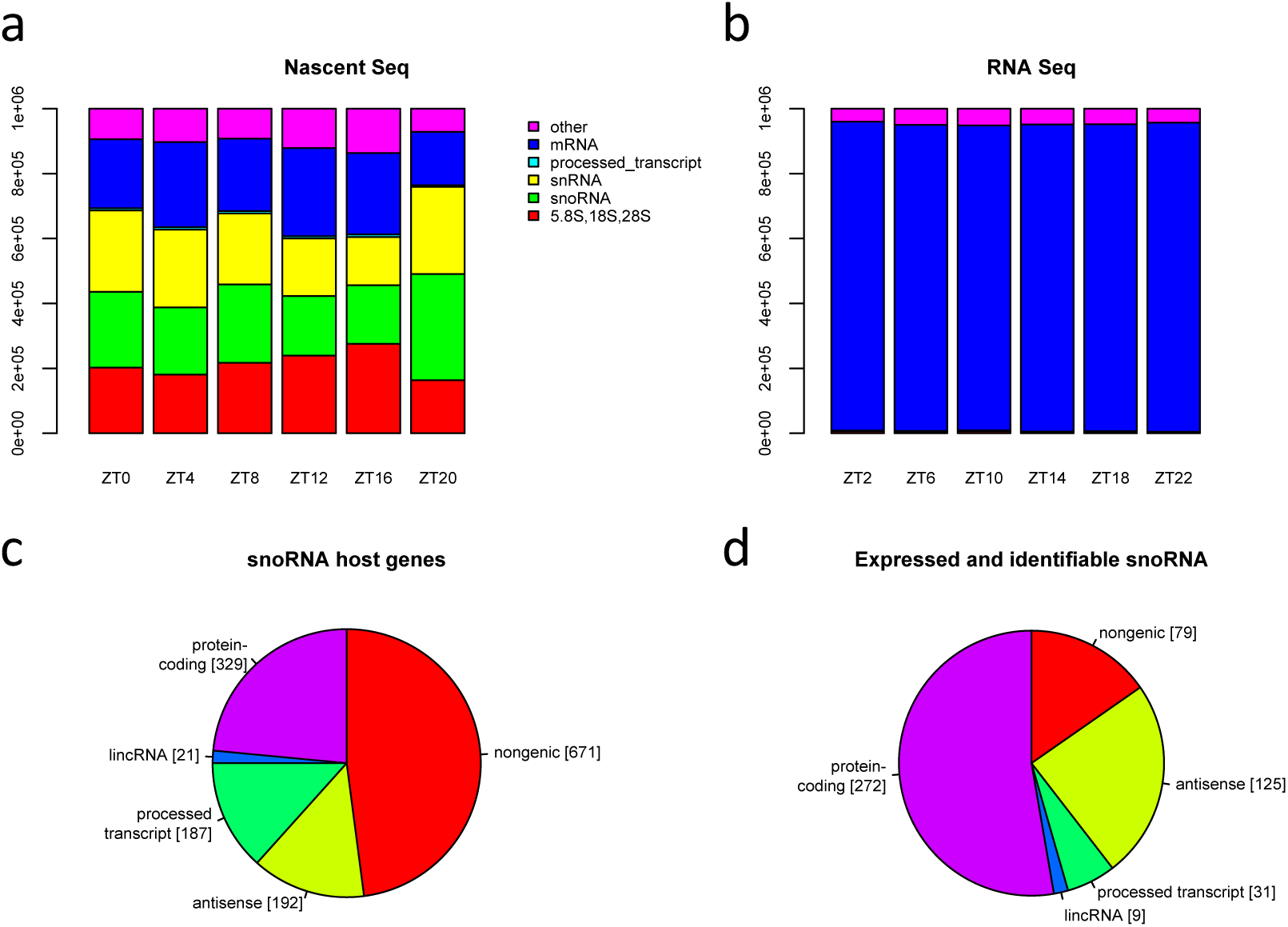
All major categories of non coding RNA are captured by nascent sequencing. **a** Stacked bar charts show the total expression of five selected RNA biotypes in nascent sequencing, and in RNA sequencing data **b**, at 6 time points. Quantification is in TPM and hence sums to 10^6^ at each time point. **c** Chart depicts the numbers of small nucleolar RNAs annotated in Ensembl classified according to host gene biotype, designated antisense if on the opposite strand to an overlapping gene, else designated nongenic. **d** The number of small nucleolar RNAs categorised as in **c** that are both expressed and identifiable by a uniquely mapping read in nascent sequencing data.

The categorisation of all Ensembl snoRNAs according to the gene type of the host gene (if any), and the numbers expressed in nascent sequencing are indicated in Figs 1c and 1d. Overall, we found 516 snoRNA (37%) to be both expressed and identifiable in mouse liver. For snoRNA hosted by protein-coding genes the fraction rises to 83%. A smaller proportion of antisense snoRNA were expressed and identifiable, but the fraction was still surprisingly high at 65%. In contrast, only 12% of other nongenic snoRNA meet these criteria. As this analysis was based on the alignment of reads to sequences, duplicate sequences had to be eliminated (101 snoRNA genes had one or more duplicates and were replaced by 17 exemplars to give a total of 1400 unique genes). Supplementary File 1 lists the snoRNA in mm10 along with their locus, that of their host, RFam family, snoRNA type, equivalence class and whether expressed or not.

We then examined whether nongenic snoRNA tend to have higher sequence similarity with other snoRNA as an explanation of their lack of identifiability. Using Blast, we built sets of genes with sequence alignments from 85% to 100% and found that snoRNA with processed transcript hosts were more prevalent than expected in the 85% similarity set and that nongenic snoRNA were not (Fig S1). The number of identifiable genes (those with uniquely-mapping reads) reduced as sequence similarity increased, and in the 85% similarity set anti-sense snoRNA were more identifiable than expected and nongenic snoRNA less so (Fig S1). However, at 85% similarity, only 71 nongenic genes were not identifiable, and so sequence similarity appeared to be only a small factor in the eight fold reduction in the number of nongenic snoRNA that were actually expressed and identifiable: This class of snoRNA does not appear to be active in mouse liver.

**Figure 2:**
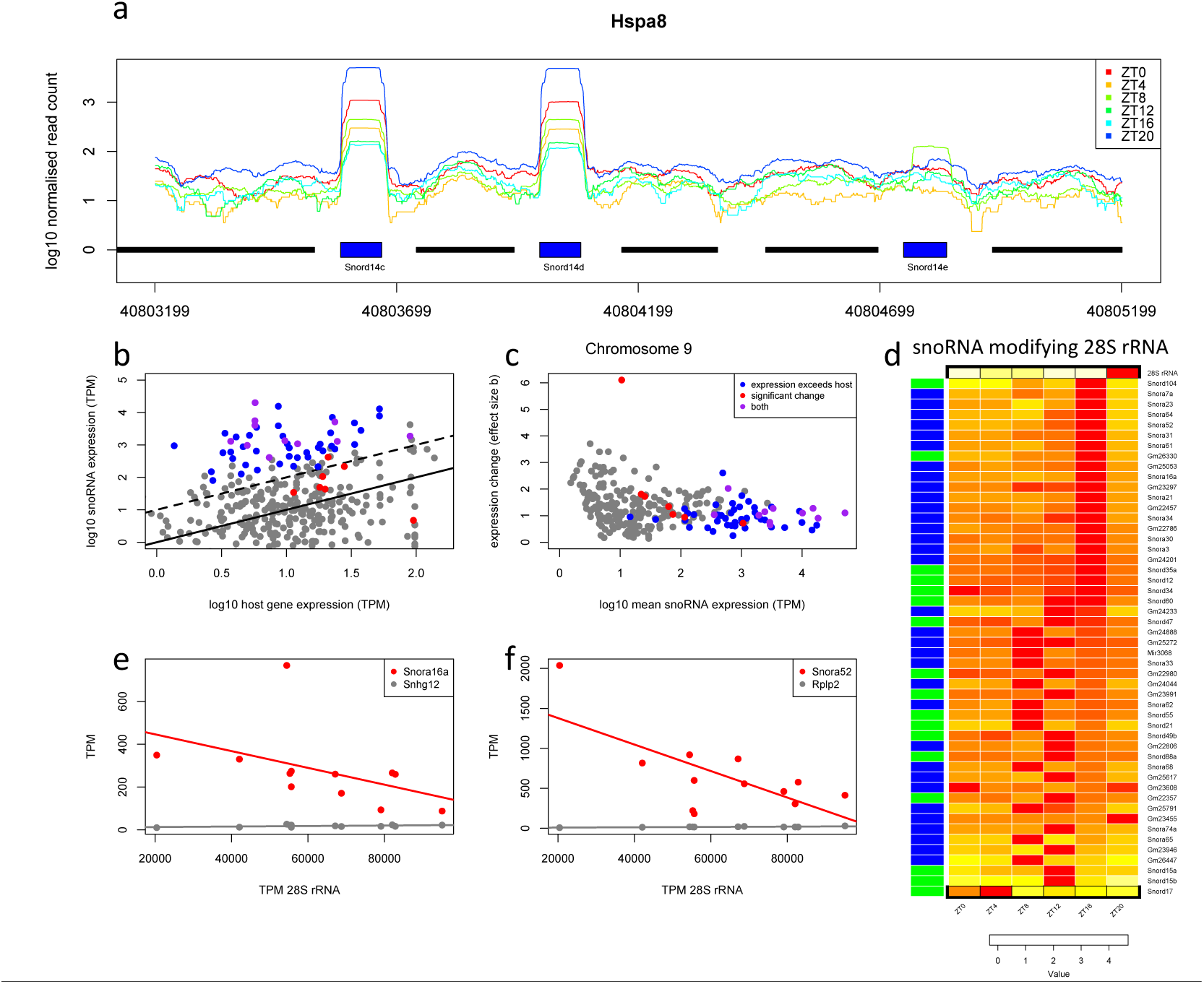
SnoRNA expression varies considerably over time. **a** Normalised read depth in nascent sequencing data over the Hspa8 locus at 6 time points. The locations of Hspa8 exons are shown by black bars, snoRNA by blue bars. Nascent sequencing depth was 70M-157M and coverage was normalised to 10^8^. **b** Scatterplot of snoRNA expression against host gene expression at ZT0. Points above the solid black line represent snoRNA with abundance greater than their host, and those above dashed black line have expression 10 times greater than their host. **c** Scatterplot of log fold change in snoRNA expression (b value calculated by sleuth) against mean expression over the time series. In **b c**, blue symbols indicate snoRNA with expression at least than ten times that of their host gene at all time points, red indicates a significant change in expression (adjusted p ≤0.05), and purple shows snoRNA satisfying both criteria. **d** Heatmap of the expression of 28S rRNA and snoRNA known to modify 28S. Scale is log 10 difference in TPM from minimum. 28S rRNA (top row) has peak expression at ZT12-ZT16 whereas snoRNA known to interact with 28S have minimum expression at this time. Box H/ACA snoRNA are indicated by blue side colours and box C/D by green. **e f** Scatterplots of the expression of selected 28S-modifying snoRNA against 28S expression for the 12 samples available (2 replicates at six time points). **e** Scatterplot of Snord92 (host gene Wdr43) and **f** Snora52 (host gene Rplp2) against 28S expression, lines show linear regressions for snoRNA (red) and host gene (grey).

### 2.3 A subpopulation of snoRNA have time-varying expression greatly in excess of their host gene

The difference in expression between genic snoRNA and their host genes was a striking feature of the nascent sequencing data. For example, Snord14c and Snord14d were 20 times more highly expressed than their host Hspa8 at certain times of the day (Figs 2a and S2). Many snoRNA were consistently more highly expressed than their nascent host gene, indeed, 56 were at least 10 times more greatly expressed than their host gene at all time points (Fig 2b). To assess the change in expression in these genes we might consider the fold change between maximum and minimum values over the time course. However, simply requiring a threshold of a 2-fold change in mean expression would lead to the conclusions that 63% of all snoRNA with a host showed differential expression, and that a comparable fraction (66%) of snoRNA with expression in excess of their host were differentially expressed. In fact, testing for differential expression using the Wald test (implemented in sleuth [20]) such that variability between replicates is accounted for led to a very different conclusion: 4% of snoRNA with a host (21 genes) showed significant changes and 25% of snoRNA with expression in excess of their host were differentially expressed. The 21 genes identified had adjusted p values ≤0.05 after accounting for the testing of 98327 transcripts, and the same set were significant if we considered only snoRNA and chose a conservative threshold of 0.005 after Benjamini Hochberg correction (a conservative threshold is warranted to account for the selection of minimum and maximum values over the time course). The extent and significance of the fold changes in snoRNA expression over the day are indicated in Fig 2c by the plot of e ect size (the b value computed by sleuth, proportional to log fold change) over the time course against mean expression. Known modifiers of 28S are among the 14 genes satisfying both criteria in Figs 2b and 2c: Snord17, Snora23, Snora65, Snora74a, and Gm23946. (Supplementary File 2 lists these genes and provides their expression data.) These properties of snoRNA abundance raise questions as to the relationship between snoRNA and the rRNA they modify, and raise the possibility that some snoRNA may be cyclically expressed. The limitations of assessing circadian regulation through comparisons of maximal and minimal expression are also evident and we address these below.

### 2.4 The expression of snoRNA known to modify 18S and 28S rRNA is negatively correlated with rRNA expression

To obtain a reliable functional annotation of snoRNA, we found exact sequence matches for Ensembl genes in the snOPY database [18] and thereby accessed curated data on the modification of rRNA by snoRNA. This resource also provided informative names for many mouse genes whose names in Ensembl begin ‘Gm’ (following snOPY usage, these names are capitalised). Utilising this information, we observed many snoRNA known to modify 18S and 28S rRNA to have minimum expression at ZT12 or ZT16, that is, precisely the time when 18S and 28S expression reached a peak (Figs 2d and S3) and to increase thereafter. To quantify this unexpected relationship we derived linear models for the expression of each snoRNA as a function of 28S expression, and similarly for the host genes of these snoRNA and 28S expression. The scatterplots of Figs 2e and 2f illustrate two examples where snoRNA expression is negatively correlated with 28S and the host gene is positively correlated with 28S. To assess the statistical significance of these correlations, we compared the number of snoRNA targeting 28S that are negatively correlated with 28S with the numbers negatively correlated in the remainder of expressed genes at a specified value of *R*^2^ using the hypergeometric test (and similarly with positively correlated genes, and for host genes). Rather than select a value of *R*^2^ apriori, we assessed overrepresentation for *R*^2^ from 0 to 1, and found the negative correlation of snoRNA to be significant up to an *R*^2^ of 0.56 (p=0.009). The fractions of snoRNA and host genes with positive and negative correlations to 28S are plotted in Fig S4 where it can be seen that as *R*^2^ increases the number of genes reaching this level of correlation reduces until there are insu cient genes to test. A similar pattern is found for snoRNA modifying 18S. It should be noted that to counteract the variation in total rRNA, rRNA genes were removed and the expression of other genes rescaled to 10^6^ in the above analysis. This analysis was repeated by quantifying counts of uniquely mapping reads (see Methods) and again we saw a striking increase in counts at ZT20 in comparison with ZT16 (Fig S5).

**Figure 3:**
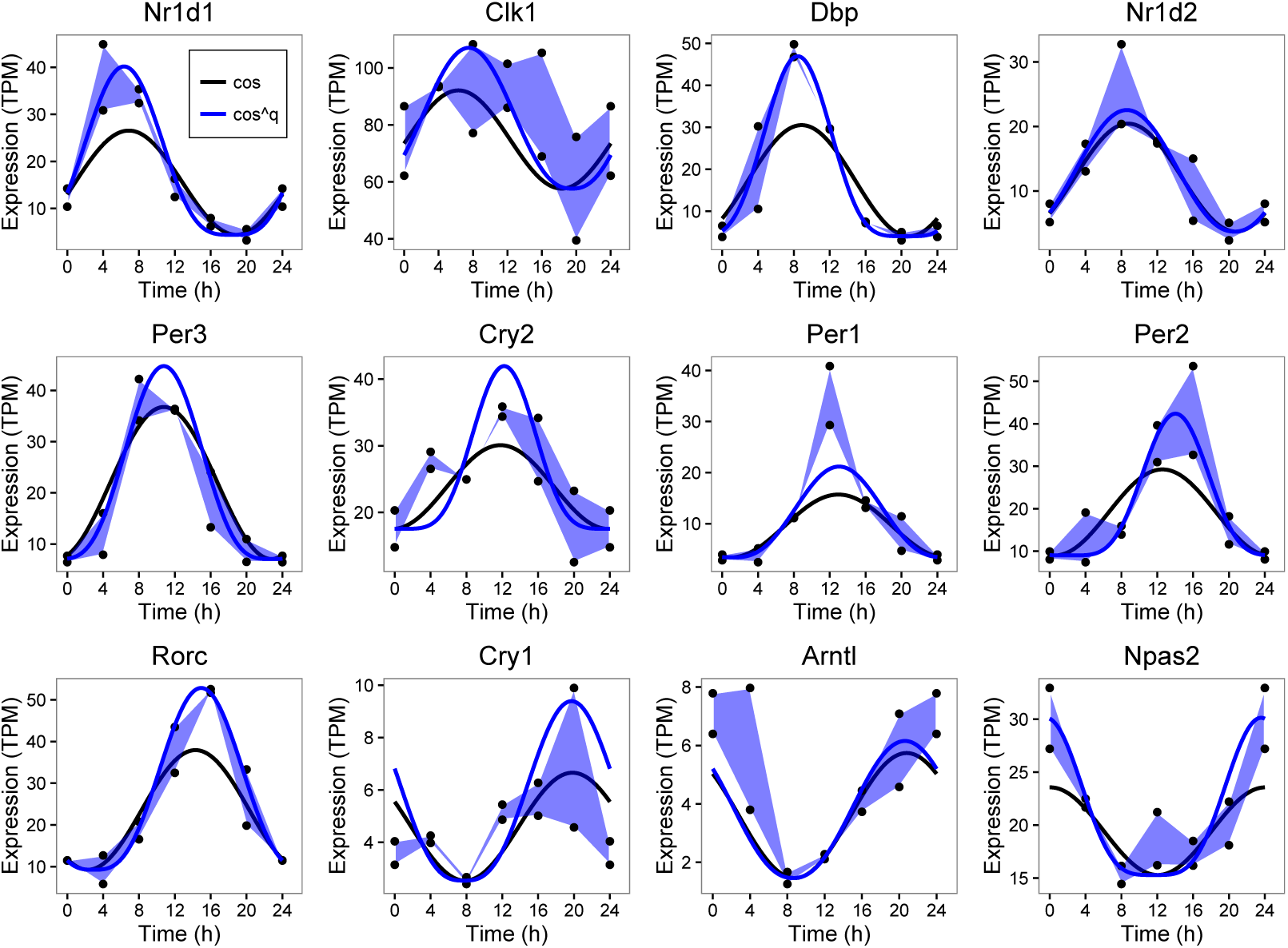
A novel cosine model better fits nascent seq data for known clock genes. Expression of twelve established clock genes plotted as a polygons (blue shaded areas) between the maximum replicate data values across the time series and their minimum. Black symbols are data values. The solid lines are the best fit of the standard cosine model (black) and the cosine^*q*^ model (blue) to the median of the replicates at each time point.

A positive correlation between nascent mRNA and pre-rRNA potentially reflects coordinated transcriptional regulation as reported for ribosomal protein genes [8]. In contrast, the negative correlation between snoRNA and pre-rRNA expression implicates post-transcriptional mechanisms that may include intra-nuclear tra cking and release from the ribosome precursor.

### 2.5 Inference of circadian rhythms: A novel method combining residual error and standard deviation of phase

To further analyse potential rhythmic oscillations in snoRNA and host genes, we adopted an established false discovery method based on Fourier analysis named F24 [21] (as used in [5]) as an initial filter. Genes with p value for their F24 statistic of greater than 0.2 were not considered further. Exploring alternative mathematical models of circadian dynamics we found that nascent expression data was better fitted by a cosine function raised to a power, creating a more peaked cycle, than a simple cosine function.

The improvement in fit of the new model in comparison with the standard cosine model for twelve clock genes is illustrated in Fig 3. When assessing the goodness of fit of circadian models to data we found it important to account for the variability between replicates both in the computational analysis and in visualisation. The variability in expression between replicates across the time series is readily perceived by plotting curves between the upper values across the time series, and similarly between the lower values, forming a polygon (as in Fig 3). The cosine models were fitted to the median of the replicates at each time point and so the cosine curve would ideally be equally spaced between the upper and lower replicates at each time point. Recall that each data point is from a different mouse, hence the importance of accounting for biological variability.

This modified cosine model with period 24 hrs, with period 12 hrs and a linear model were fitted to each time series to assess the fit of a true circadian rhythm, a rapidly oscillating signal (likely noise given the sampling frequency of this data, but potentially due to transcription factor binding [22]) and a gradual change in expression respectively. The Bayesian evidence for each of the three models was calculated using nested sampling [23, 24] and time courses were designated circadian where the evidence for the 24 hr cycle was ten times that for the alternative models. The nested sampling algorithm infers the phase and its standard deviation, both of which are of interest in assessing rhythmic behaviour. The likelihood function accounts for the consistency between replicate data, giving less weight to times where replicates di er more (see Methods).

In line with comparable methods, 9% of protein coding genes were found to be circadian. To compare the results of our method with published results in more detail, the phase calculated by nested sampling is plotted against the phase calculated by the Fourier method in Fig S6 (a) for protein-coding genes designated circadian in [5] (*R*^2^=0.53, p≤2e-16). To further refine the set of circadian genes, those whose phase could not be inferred accurately, or whose fit to the cosine model was less good (as determined by the standard deviation of the phase and the residual (L1) error respectively, see Methods) were excluded. As these two measures can be traded off, we defined a radial score that combines them, and excluded the worst scoring 5% of these circadian genes (Fig S6 (b)). The distribution of phase values by our method and by the published method (where both the quantification of expression and phase calculation di er) are comparable (Fig S6 (c)). The range of values chosen for the power parameter (*q*) in the proposed cosine model is shown in Fig S6 (d). Values of *q* greater than 1, the value of the standard model, were chosen extensively. Plots of nascent and RNA sequencing data and the fitted models for 12 clock genes can be found in Fig S7. Turning to snoRNA and their host genes, the filtering and selection procedure yielded 43 circadian snoRNA and 26 circadian host genes (Fig S6 (e)). The absolute radial score threshold determined from circadian genes was also applied in this case.

### 2.6 A subpopulation of snoRNA show cyclical expression

Thirteen snoRNA located in introns were found to be cyclically expressed, including Snord35b, Snord57 and Snord14d. The peak expression of these snoRNA occurred across the day with some preference for the beginning or end of the day (Fig 4a). Thirty nongenic snoRNA were cyclically expressed, showing peak expression within a more defined period 4-16hrs after dawn (Fig 4b). The distribution of phase values (Fig 4c) illustrates the di ering peak times of these two populations of snoRNA. We next looked for cyclically expressed host genes in both nascent sequencing and RNA sequencing data and identified 26 and 14 cyclic host genes respectively (Fig S8). Of the 30 snoRNA whose host showed cyclic expression in nascent seq data, two were found to be cyclically expressed and we observed one of these to be in anti-phase with its host and the other to be in phase (Figs S9 and S10). Thus we found only minimal overlap between snoRNA and host expression patterns possibly indicating that their cyclic behaviour is regulated by mechanisms in addition to transcriptional activation. The model parameters for cyclical snoRNA and their host genes can be found in Supplementary File 3.

As is apparent from Fig 4, few of the cyclical snoRNA are currently designated ‘Snora’ or ‘Snord’ which indicates a lack of recognition of their status in mouse. However, from the snOPY database we identified SNORA21 (Gm25821), SNORA46 (Gm26493), SNORD88 (Gm26247), SNORD115 (Gm26337), and three SNORA17 genes (Gm25272, Gm24607, and Gm24656) among the cyclic snoRNA with host genes. Considering cyclic snoRNA without host genes, we identified SNORA63 (Gm23679), SNORA71 (Gm22797), SNORD86 (Gm23706) and seven SNORA17 genes (Gm22778, Gm26421, Gm23910, Gm24375, Gm24556, Gm23674 and Gm22670). Of note, genes in the SNORD88 and SNORD115 families are associated with the regulation of splicing [10].

**Figure 4:**
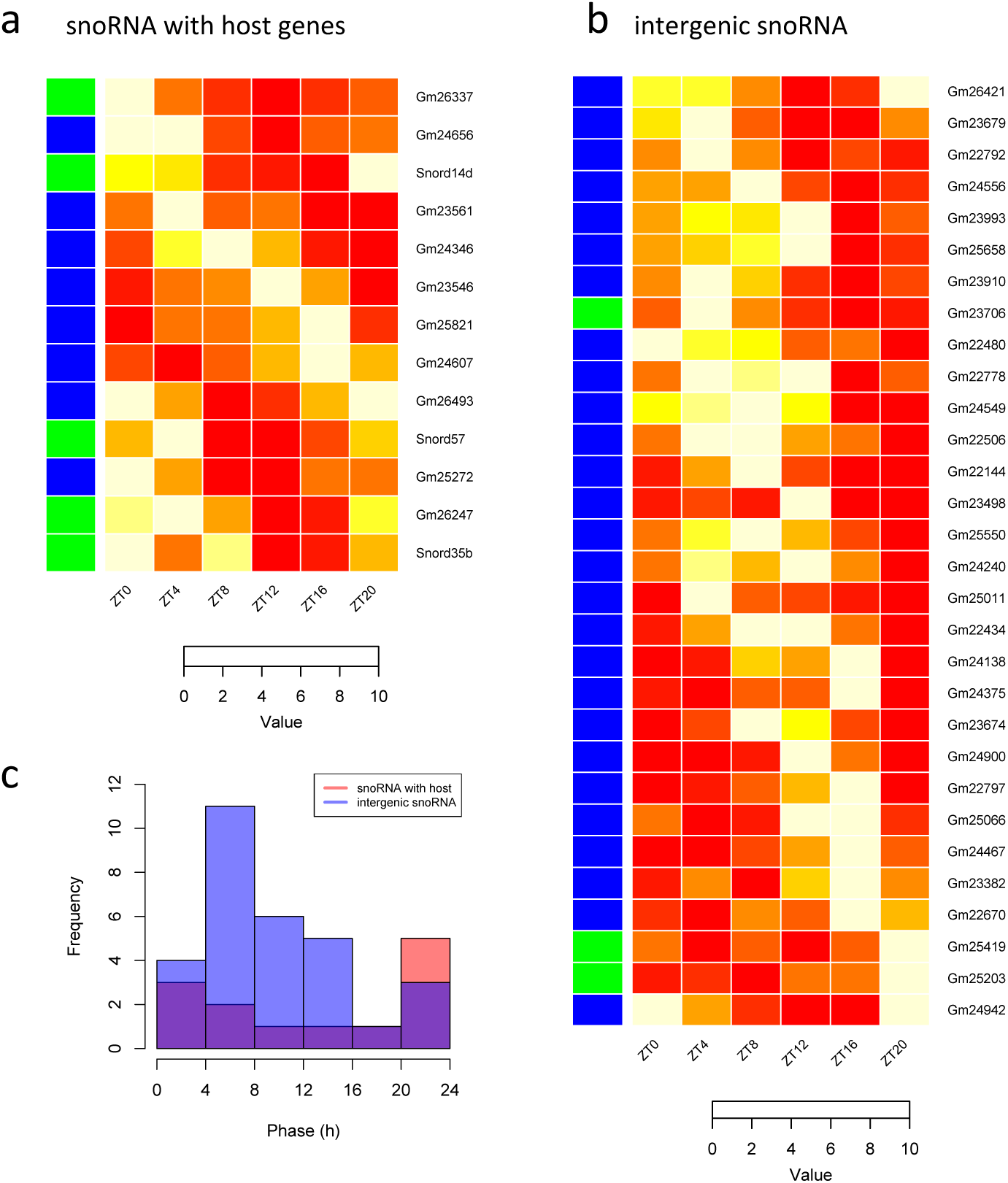
Cyclically expressed snoRNAs. **a** Heatmap of the expression of 13 snoRNAs with host genes that are inferred to be cyclic. **b** Heatmap of the expression of 30 cyclic intergenic snoRNAs. Heatmap rows are ordered by the inferred phase of the cosine function, box H/ACA snoRNA are indicated by blue side colours, and box C/D by green. Expression is scaled to range from 0 to 10 (see Methods). **c** Histogram of the phase of snoRNA in **a** and in **b**.

The abundance of cyclic snoRNA was on average 1.5 times that of their host genes. Of these genes, only SNORA46 (Gm26493) was among the set of snoRNA with consistently high ratios of expression relative to their host (at least ten times greater). None of the cyclic snoRNA were among those found to have statistically significant fold changes (Fig 2b) thus these populations of snoRNA were disjoint.

**Figure 5:**
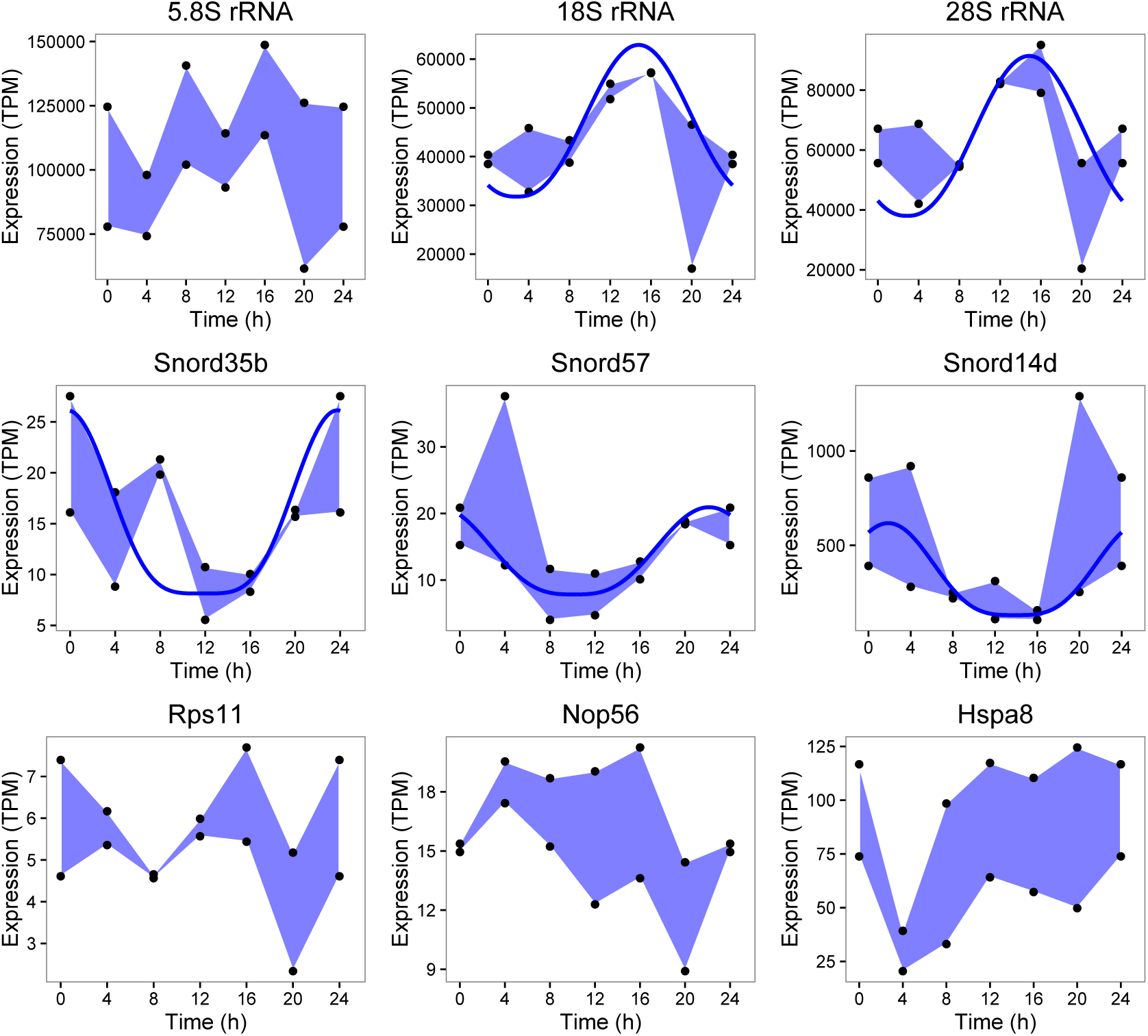
Ribosomal RNA, snoRNA and host gene expression. Expression of 5.8S, 18S and 28S rRNA (top row), selected cyclic snoRNAs (middle row) and their respective host genes (bottom row; Rps11 hosts Snord35b, Nop56 hosts Snord57 and Hspa8 hosts Snord14d) plotted as polygons (blue shaded areas) between the maximum replicate data values across the time series and their minimum. Where expression was inferred to be cyclic, the best fitting cosine^*q*^ model is indicated by a solid blue line. Notably, Snord57 modifies 18S and the protein of its host Nop56 is a component of the box C/D ribonucleoprotein complex.

### 2.7 18S and 28S rRNA are cyclically expressed

Applying the circadian modelling introduced above, we next determined that the temporal variations noted earlier in both 18S and 28S rRNA were indeed circadian, while 5.8S expression dynamics did not pass the initial false discovery filtering step. The cyclical patterns of these transcripts are shown in Fig 5 along with selected circadian snoRNAs (Snord35b, Snord57 and Snord14d; middle row) and their respective host genes (bottom row). Snord57 and Snord14d are known to modify 18S rRNA and it is readily seen that their expression profiles show starkly contrasting phase.

Of the 10 cyclic snoRNA with host genes that had matches in the snOPY database, four modify 28S: Snord35b, SNORA21, SNORA17 (Gm25272) and SNORD88. Three modifiers of 18S were found among the cyclic snoRNA with hosts: Snord57, Snord14d, and SNORA46. Thus, we found the majority of cyclic snoRNA with host genes to be associated with rRNA modification, however as the majority of these genes modify 28S or 18S this number did not constitute a statistical enrichment. The cyclic snoRNA without hosts we found included SNORA17 (Gm24375) a modifier of 28S, and SNORA71, a modifier of 18S.

### 2.8 Highly-expressed nongenic snoRNA have a distinct chromatin signature

Histone modifications H3K4me3 and H3K27ac have been shown to vary rhythmically around gene promoters in mouse liver [14], and the rhythmic recruitment of PolII at the promoter has been demonstrated to oscillate in phase with RNA polymerase II (PolII) levels on the gene body indicating that it is the recruitment of PolII rather than its release that is critical to diurnal transcription [13]. A set of strong circadian promoters has been proposed to drive circadian genes with high amplitude and high average expression, and is associated with high paused PolII levels (relative to H3K4me3) and the extension of H3K4me3 into the gene body [25].

To investigate whether nongenic snoRNA have a chromatin signature that might support their transcription as independent genes, and to explore any temporal variations indicative of circadian expression, we mapped the PolII and H3K4me3 time series data from mouse liver published by Le Martelot [13] and located peaks at each time point, and in the combined data using MACS2 [26]. The abundance of PolII and H3K4me3 around clock gene promoters, and the variation in these signals is shown in Fig S11 for Per2 and Nr1d1. Consistent with previous studies, a substantial peak in PolII was observed at the gene start with peaks in H3K4me3 downstream. Of the twelve clock genes examined, PolII levels decreased towards background levels at one or more time points in three cases (Per2, Dbp, and Npas2) whereas H3K4me3 levels remained above background across the day in all cases. We then examined the chromatin signature of three snoRNA known to be independently transcribed, namely, Rnu3a (U3), Snord13 and Snord118 [19, 27], and found a distinctive peak in PolII at the gene start in all three cases (Figs 6 and S12). A considerable temporal variation in this signal was also apparent. These genes overlapped with peaks in PolII and H3K4me3 called by MACS2 and so we searched for other nongenic snoRNA that shared these properties and found six: Snord104 and SNORA76 (Gm22711) (which are clustered as in human [19]), Snora57 (reported to be monocistronic in [18]), Snora17, Gm25501, and Gm23596 (which are antisense to Ank2 and intergenic respectively). In the cases of Rnu3a and Snord13, the upstream peaks in H3K4me3 were over the start of an adjacent gene on the opposite strand (Gtf3c6 and Tti2 respectively). Although Snora17 has no annotated host gene in the release of Ensembl we have adopted, it overlaps Snhg7 in Refseq. The major peaks in the chromatin signals around Snora17 were located over the Refseq host gene start (with minor peaks over the gene itself) which support the existence of the host.

The eight snoRNA we characterise as independently transcribed had higher PolII, H3K4me3 and nascent seq expression than did nongenic snoRNA that lack overlapping MACS2 peaks in PolII and H3K4me3 (p≤2.7e-4 by Wilcoxon test). The input PolII and H3K4me3 levels of these genes did not differ from that of the remaining nongenic snoRNA (to determine an overlap the snoRNA gene locus was extended by 200 bases, and the expression of these extended features was quantified in RPKM). It is readily evident in Fig S13 that these eight genes form a distinct cluster of highly expressed snoRNA with corresponding chromatin marks. In addition we found Snord60 and Snora78 (which overlap short antisense transcripts Rab26, and Snhg9 respectively) to have similar chromatin signatures.

The PolII signal of each of the eight independently transcribed snoRNA had a distinct minimum at ZT6, and for all except Rnu3a there was a dip in nascent seq expression at ZT8 relative to ZT4, followed by an increase at ZT12 (Fig S14) and variable expression thereafter. The di ering sampling times of these data made the assessment of any correlations unreliable. The minimum PolII signal was at least twice the background, and the log2 fold change of the maximum signal (relative to the same background) was at least 1.4 greater than the minimum which again indicated a notable temporal variation. Snord13 was the most circadian with a F24 FDR 0.12 (three other genes also had p values less than 0.2). Our Bayesian method could not be applied to the chromatin data as there were no replicates. None of these snoRNA had cyclic expression in the nascent seq data. However, the log2 fold change in nascent expression was in the range 0.9-2.9 when maximum and minimum expression over the day were compared, and therefore temporal variation was evident in all cases. The H3K4me3 signal dipped at ZT14 or ZT18 in six cases but with less pronounced fold changes over background than for PolII (Fig S14). Applying the F24 FDR test, we found three snoRNA to have p values for H3K4me3 expression less than 0.2 (Rnu3a, Snord104 and Gm22711).

**Figure 6:**
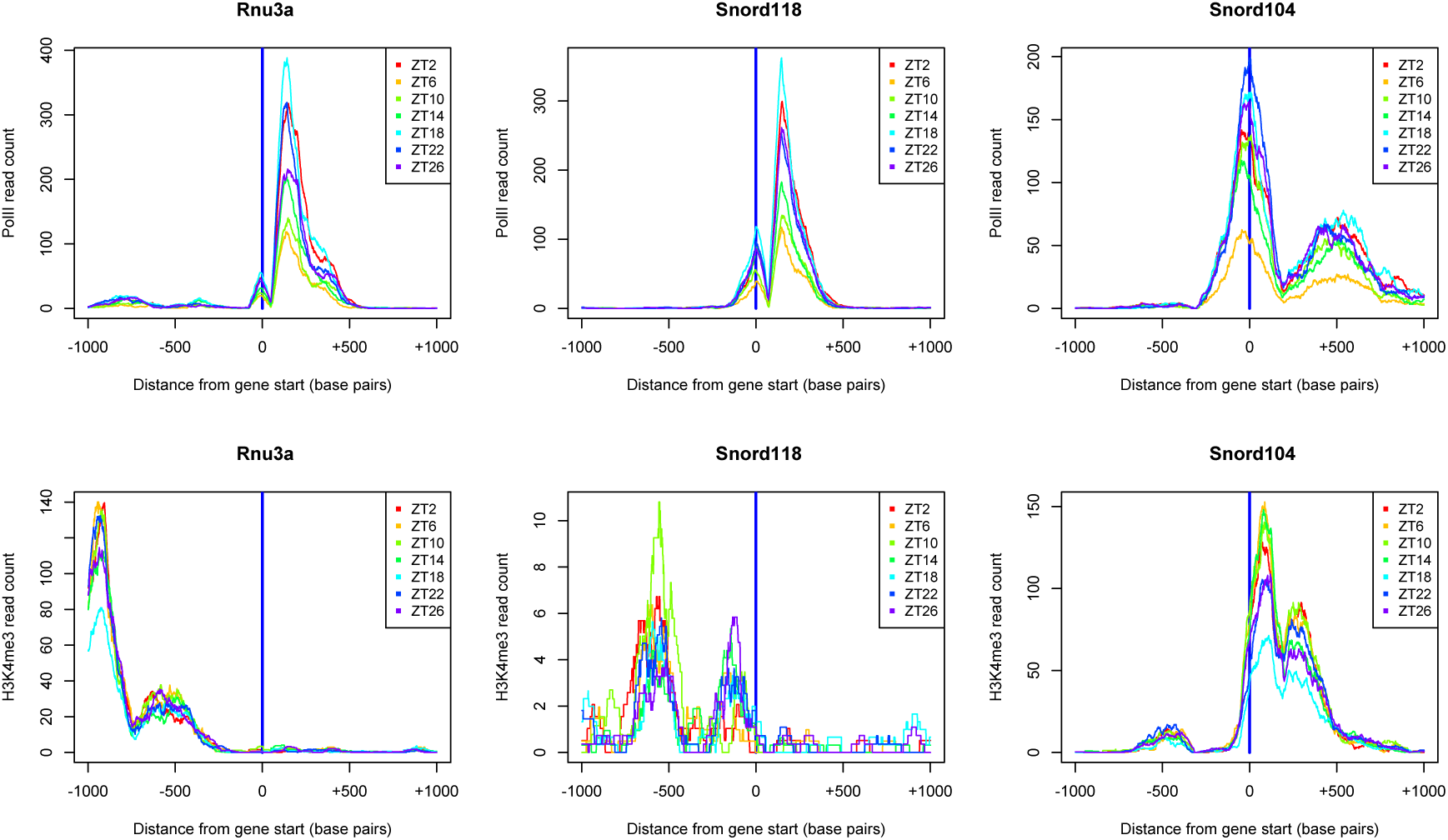
Independently transcribed snoRNA have a distinct chromatin signature. Normalised read depth in a 2kb region centered on the snoRNA gene start, and oriented in the direction of transcription, is shown for selected independently transcribed snoRNA for RNA Polymerase II (top row) and for H3K4me3 ChIP sequencing data (bottom row) data at seven time points. A peak in PolII over the gene and an adjacent peak in H3K4me3 are characteristic chromatin features.

## 3 Discussion

Using novel computational statistical techniques we have uncovered previously unrecognised patterns in the abundance of nuclear pre-rRNAs and snoRNAs, and correlations between them. A population of snoRNA that were at least ten times as abundant as their nascent host gene, some with statistically significant diurnally-varying (but not cyclical) expression was identified. The expression of snoRNA that modify 18S and 28S was typically in antiphase with that of the target rRNA precursor, as evidenced by negative correlations in abundance.

We found the expression of ribosome precursors 18S and 28S rRNA to follow a circadian rhythm in mammalian liver, peaking at ZT16, and that snoRNA including Snord14d, Snord35b, and Snord57 also had cyclical expression patterns in this tissue. Snord57 is known to modify 18S and the protein of its host gene, Nop56, is a component of the box C/D ribonucleoprotein complex. Thus there may be common underlying regulation that we are now beginning to unravel. The scope for confirmation of our findings in other time course data was limited as gene expression is typically measured by microarray, or by poly A+ and rRNA depleted RNA sequencing. However, a small number of microarray probes in [8] did match snoRNA and the expression of three cyclical snoRNA was reproduced (Fig S15).

The intersection between circadian snoRNAs and circadian host genes was minimal as only two cases were found. In the first, snoRNA and host expression were in antiphase, in the second, expression was in phase. Given the overall proportions of cyclic genes in these categories, there was no enrichment for cyclic host genes among cyclic snoRNAs, hence no evidence for cyclic transcription as the key regulator. As for messenger RNA [2, 5], mechanisms in addition to transcription must contribute the regulation of cyclic nuclear snoRNAs.

The correlation of snoRNA host gene and rRNA expression may be the result of co-regulation with rRNA as reported for ribosomal protein genes [8]. An antiphase relationship between many snoRNA and their pre-rRNA target is more surprising, and may show an upregulation of snoRNA biogenesis in anticipation of the increased rRNA levels that peak around ZT16, or may be due to a release (or relocation) of snoRNA from the previous cycle of rRNA maturation that restores their abundance in the nucleus. The rapid recovery in the expression of many snoRNA seen at ZT20 (Figs 1d, S3 and S5) argue for the latter.

Mature snoRNA are concentrated in the nucleolus, however they undergo extensive intranuclear tra cking during biogenesis [28]. Indeed, the box C/D motif functions as the nucleolar localisation signal [29]. In addition, snoRNAs have been found to be involved in splicing outside of the nucleolus [30]. Human U8 (SNORD118) snoRNA precursors have been found in cytoplasmic extracts in levels comparable with those in nuclear extracts [31] but this does not appear to be a typical biogenesis pathway [32]. Thus for a number of snoRNA, variation in abundance may be attributed in part to cytoplasmic tra cking, and possibly to tra cking between nuclear structures, as well as to their established role in rRNA biogenesis.

Little is known about the role of the chromatin environment as a potential regulator of independently transcribed snoRNA. We found peaks in RNA polymerase II over the gene locus and adjacent peaks in H3K4me3 to be signatures of independently transcribed snoRNA, and, in addition, mean PolII and H3K4me3 levels correlated with mean snoRNA transcript abundance. Time-varying but noncyclic patterns were found in these chromatin marks, with a distinct dip in PolII at ZT6 that may indicate a common regulatory input for this class of snoRNA.

Differences in phase of clock-regulated genes in different organs have been reported [3, 11, 33], o ering insights into the coordination of the peripheral clocks. Our methodology is particularly suited to such investigations as it yields standard deviations for key model parameters such as phase, and the potential to model multiple data sets in an integrative manner.

## Materials and methods

### Definition of cosine models

Circadian rhythms were modelled by a cosine function that varied between 0 and the maximum expression *a*, with peak expression (i.e. phase) *p* minutes after time 0, raised to the power *q* (Eq 1).

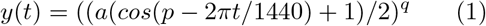

Parameters *a*, *p* and *q* were constrained by the following prior ranges:

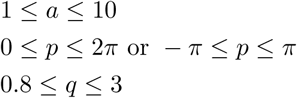

All time series data were scaled such that the minimum median value was 0 and the maximum median was 10, hence *a* could be at most 10. Two alternative constraints on *p* were used to ensure that the fitted value of this parameter did not lie at the end of the prior range. This might occur for *p* close to 0 or 2π in which case the alternative prior centered on 0 (2π) was used − π ≤ *p* ≤ π.

The fit between the cosine models and expression data was assessed using the nested sampling algorithm to calculate the log of Bayesian evidence (also known as the marginal likelihood), log Z [23] from the likelihood function and the prior. All priors were selected uniformly from a range bounded by maximum and minimum values given above. A likelihood based on the l1-norm was defined by Eqs. 2 and 3 [34]. Equation 2 defines the normalising constant *ϵ_t_* as the expected value of the moduli of the difference between the replicate observations at time *t* (*x_t_*) and the value predicted by the kinetic model (*µ_t_*). The product of the probabilities of the median observation at time 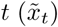 defines the likelihood function for a time series *x* of *m* samples (Eq. 3). Maximisation of this likelihood minimises the sum of the moduli of the residuals (rather than their squares) on the basis that the testable information is restricted to the expected value of the modulus of the difference between theory and experiment. Should we know both the mean and variance, maximum entropy considerations would lead instead to the Gaussian distribution [34].

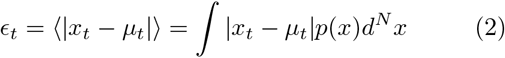

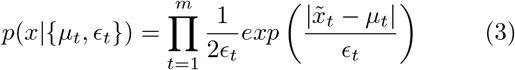

Bayesian evidence values and model parameter estimates (and their standard deviations) were computed using nested sampling for each time series that passed an initial FDR test (the F24 test [21] with p≤0.2). A cosine model with a 12 hour period and a linear model were also fitted to each time series. Time series where the log Z for the 24 hr cosine model was ten times greater than that for the alternative models were considered circadian if, in addition, they passed a test on the standard deviation of phase and L1 error. The threshold for the final radial score test was derived empirically from genes found to be circadian in earlier studies [5]. R code for nested sampling is provided in Supplementary File 4.

### Processing of sequencing data

The gene annotation file for GRCm38 was downloaded from Ensembl (version 84) and processed with bedtools [35] and in R to identify snoRNA, their locus, snoRNA host genes and their locus, and gene biotypes. SnoRNA - host gene assignments were reviewed manually using the IGV genome browser. Additional data on RFam families was downloaded from the EBI, and data from the snoPY database [18] was also utilised.

Nascent and RNA sequencing time series data was downloaded from GEO GSE36916 [5]. Coding and non-coding transcripts for mouse genome GRCm38 were downloaded from Ensembl to which the 5.8S, 18S and 28S pre-rRNA sequences were added from [18] to create an index file for quantification in TPM using Kallisto [17]. TPM values for genes were summed from those of their transcripts. Reads were also mapped to Ensembl GRCm38 using bowtie2 (using parameters -L 18 -N 1 -k 20; Fig S2) [36]. Uniquely-mapping reads were extracted using samtools [37], and unique read counts for snoRNA genes found using htseq-count [38]. These counts were used to determine snoRNA identifiability. Read pileups (Fig 2) were created from multiply-mapped reads using bedtools with output files subsequently processed in R.

Following [5], the F24 test [21] was applied to the Nascent and RNA time series data by concatenating first and second replicates to create a series from ZT0 to ZT44. We constructed the replicated time series in the same manner in order to have a sample at ZT24 while not duplicating the ZT24 sample alone (Figs S3 and S5 show ZT0-ZT24 only).

PolII, H3K4me3 and input time series data was downloaded from GEO GSE35790 [13]. Reads were mapped to Ensembl GRCm38 using bowtie2 (using parameters -k 2) and uniquely-mapping reads were extracted using samtools. MACS2 [26] was used to find peaks in uniquely-mapping PolII and H3K4me3 reads at each time point, and in the combined data. Peaks found in the combined data appeared most robust and were intersected with snoRNA locus using bedtools. Read counts and pileups for genomic features were obtained using bedtools and output files were subsequently processed in R (Figs S6, S9, S10 and S11).

## Acknowledgements

### Authors contributions

SA and CAS designed the study. SA performed the computational analysis. SA and CAS wrote the manuscript. All authors gave final approval for publication.

### Competing interests

We declare we have no competing interests.

### Funding

SA and CAS were funded by the MRC Human Genetics Unit core grant.

### Acknowledgments

We acknowledge the assistance of Prof. Naoya Kenmochi of the Frontier Science Research Center, University of Miyazaki, Japan, with the functional annotation of mouse snoRNA sequences in the snOPY database.

## References

[1] Staiger, D., Shin, J., Johansson, M., and Davis, S. J. (2013) The circadian clock goes genomic. Genome Biology, 14, 208.

[2] Benegiamo, G., Brown, S. A., and Panda, S. (2016) RNA dynamics in the control of circadian rhythm. Yeo, G. W. (ed.), RNA Processing, pp. 107–122, Springer.

[3] Yan, J., Wang, H., Liu, Y., and Shao, C. (2008) Analysis of gene regulatory networks in the mammalian circadian rhythm. PLoS Computational Biology, 4, e1000193–.

[4] Rey, G., Cesbron, F., Rougemont, J., Reinke, H., Brunner, M., and Naef, F. (2011) Genome-wide and phase-specific DNA-binding rhythms of BMAL1 control circadian output functions in mouse liver. PLoS Biology, 9, e1000595–.

[5] Menet, J. S., Rodriguez, J., Abruzzi, K. C., and Rosbash, M. (2012) Nascent-Seq reveals novel features of mouse circadian transcriptional regulation. eLife, 1, e00011–.

[6] McGlincy, N. J., Valomon, A., Chesham, J. E., Maywood, E. S., Hastings, M. H., and Ule, J. (2012) Regulation of alternative splicing by the circadian clock and food related cues. Genome Biology, 13, R54.

[7] Reddy, A. B., et al. (2006) Circadian orchestration of the hepatic proteome. Current Biology, 16, 1107–1115.

[8] Jou e, C., Cretenet, G., Symul, L., Martin, E., Atger, F., Naef, F., and Gachon, F. (2012) The circadian clock coordinates ribosome biogenesis. PLoS Biology, 11, e1001455–.

[9] Bachellerie, J.-P., Cavaillé, J., and Hüttenhofer, A. (2002) The expanding snoRNA world. Biochimie, 84, 775 – 790.

[10] Dupuis-Sandoval, F., Poirier, M., and Scott, M. S. (2015) The emerging landscape of small nucleolar RNAs in cell biology. Wiley Interdisciplinary Reviews. RNA, 6, 381–397.

[11] Zhang, R., Lahens, N. F., Ballance, H. I., Hughes, M. E., and Hogenesch, J. B. (2014) A circadian gene expression atlas in mammals: Implications for biology and medicine. Proceedings of the National Academy of Sciences of the United States of America, 111, 16219–16224.

[12] Hughes, M. E., Grant, G. R., Paquin, C., Qian, J., and Nitabach, M. N. (2012) Deep sequencing the circadian and diurnal transcriptome of drosophila brain. Genome Research, 22, 1266–1281.

[13] Le Martelot, G., et al. (2012) Genome-wide RNA polymerase II profiles and RNA accumulation reveal kinetics of transcription and associated epigenetic changes during diurnal cycles. PLoS Biol, 10, 1–16.

[14] Vollmers, C., Schmitz, R. J., Nathanson, J., Yeo, G., Ecker, J. R., and Panda, S. (2012) Circadian oscillations of protein-coding and regulatory RNAs in a highly dynamic mammalian liver epigenome. Cell metabolism, 16, 833–845.

[15] Wang, H., Fan, Z., Zhao, M., Li, J., Lu, M., Liu, W., Ying, H., Liu, M., and Yan, J. (2016) Oscillating primary transcripts harbor miRNAs with circadian functions. Scientific Reports, 6, 21598–.

[16] Makarova, J. A. and Kramerov, D. A. (2011) SNOntology: myriads of novel snornas or just a mirage? BMC Genomics, 12, 543–543.

[17] Bray, N. L., Pimentel, H., Melsted, P., and Pachter, L. (2016) Near-optimal probabilistic RNA-seq quantification. Nat Biotech, 34, 525– 527.

[18] Yoshihama, M., Nakao, A., and Kenmochi, N. (2013) snOPY: a small nucleolar RNA orthological gene database. BMC Research Notes, 6, 426–426.

[19] Dieci, G., Preti, M., and Montanini, B. (2009) Eukaryotic snoRNAs: a paradigm for gene expression flexibility. Genomics, 94, 83 – 88.

[20] Pimentel, H. J., Bray, N., Puente, S., Melsted, P., and Pachter, L. (2016) differential analysis of rna-seq incorporating quantification uncertainty. bioRxiv.

[21] Wijnen, H., Naef, F., and Young, M. W. (2005) Molecular and statistical tools for circadian transcript profiling. Methods Enzymol, 393, 341–365.

[22] Westermark, P. and Herzel, H. (2013) Mechanism for 12 hr rhythm generation by the circadian clock. Cell Reports, 3, 1228 – 1238.

[23] Aitken, S. and Akman, O. (2013) Nested sampling for parameter inference in systems biology: application to an exemplar circadian model. BMC Systems Biology, 7, 72.

[24] Aitken, S., et al. (2015) Transcriptional dynamics reveal critical roles for non-coding RNAs in the immediate-early response. PLoS Computational Biology, 11, e1004217–.

[25] Westermark, P. (2016) Linking core promoter classes to circadian transcription. PLoS Genet, 12, 1–32.

[26] Zhang, Y., et al. (2008) Model-based analysis of ChIP-Seq (MACS). Genome Biology, 9, R137.

[27] Makarova, J. A. and Kramerov, D. A. (2009) Analysis of C/D box snoRNA genes in vertebrates: The number of copies decreases in placental mammals. Genomics, 94, 11 – 19.

[28] Kiss, T., Fayet, E., Jady, B., Richard, P., and Weber, M. (2006) Biogenesis and intranuclear tra cking of human Box C/D and H/ACA RNPs. Cold Spring Harbor Symposia on Quantitative Biology, 71, 407–417.

[29] Samarsky, D. A., Fournier, M. J., Singer, R. H., and Bertrand, E. (1998) The snoRNA box C/D motif directs nucleolar targeting and also couples snoRNA synthesis and localization. The EMBO Journal, 17, 3747–3757.

[30] Falaleeva, M., Pages, A., Matuszek, Z., Hidmi, S., Agranat-Tamir, L., Korotkov, K., Nevo, Y., Eyras, E., Sperling, R., and Stamm, S. (2016) Dual function of C/D box small nucleolar RNAs in rRNA modification and alternative pre-mRNA splicing. Proceedings of the National Academy of Sciences, 113, E1625–E1634.

[31] Watkins, N. J., Lemm, I., and Lührmann, R. (2007) Involvement of nuclear import and export factors in U8 box C/D snoRNP biogenesis. Molecular and Cellular Biology, 27, 7018–7027.

[32] Pradet-Balade, B., Girard, C., Boulon, S., Paul, C., Azzag, K., Bordonné, R., Bertrand, E., and Verheggen, C. (2011) CRM1 controls the composition of nucleoplasmic pre-snorna complexes to licence them for nucleolar transport. The EMBO Journal, 30, 2205–2218.

[33] Hughey, J. J. and Butte, A. J. (2016) Differential phasing between circadian clocks in the brain and peripheral organs in humans. Journal of Biological Rhythms, 31, 588–597.

[34] Sivia, D. and Skilling, J. (2006) Data analysis: A Bayesian tutorial. OUP.

[35] Quinlan, A. R. and Hall, I. M. (2010) BEDTools: a flexible suite of utilities for comparing genomic features. Bioinformatics, 26, 841–842.

[36] Langmead, B. and Salzberg, S. L. (2012) Fast gapped-read alignment with Bowtie 2. Nature Methods, 9, 357359.

[37] Li, H., Handsaker, B., Wysoker, A., Fennell, T., Ruan, J., Homer, N., Marth, G., Abecasis, G., Durbin, R., and Subgroup,. G. P. D. P. (2009) The sequence alignment/map format and SAMtools. Bioinformatics, 25, 2078–2079.

[38] Anders, S., Pyl, P. T., and Huber, W. (2015) HTSeqa python framework to work with high-throughput sequencing data. Bioinformatics, 31, 166–169.

